# **PANOPLY:** Omics-guided drug prioritization method tailored to an individual patient

**DOI:** 10.1101/176396

**Authors:** Krishna R. Kalari, Jason P. Sinnwell, Kevin J. Thompson, Xiaojia Tang, Erin E. Carlson, Jia Yu, Peter T. Vedell, James N. Ingle, Richard M. Weinshilboum, Judy C. Boughey, Liewei Wang, Matthew P. Goetz, Vera Suman

## Abstract

**Purpose:** The majority of cancer patients receive treatments that are minimally informed by omics data. We propose a precision medicine computational framework (PANOPLY: **<u>P</u>**recision c**<u>a</u>**ncer ge**<u>no</u>**mic re**<u>p</u>**ort: single samp**<u>l</u>**e inventor**<u>y</u>**) to identify and prioritize drug targets and cancer therapy regimens.

**Methods:** The PANOPLY approach integrates clinical data with germline and somatic features obtained from multi-omics platforms, and apply machine learning, and network analysis approaches in the context of the individual patient and matched controls. The PANOPLY workflow employs four steps (i) selection of matched controls to the case of interest (ii) identification of case-specific genomic events (iii) identification of suitable drugs using the driver-gene network and random forest analyses and (iv) provide an integrated multi-omics case report of the patient with prioritization of anti-cancer drugs.

**Results:** The PANOPLY workflow can be executed on a stand-alone virtual machine and is also available for download as an R package. We applied the method to an institutional breast cancer neoadjuvant chemotherapy study which collected clinical and genomic data as well as patient-derived xenografts (PDXs) to investigate the prioritization offered by PANOPLY. In a chemotherapy-resistant PDX model, we found that that the prioritized drug, olaparib, was more effective than placebo in treating the tumor (P < 0.05). We also applied PANOPLY to in-house and publicly accessible multi-omics tumor datasets with therapeutic response or survival data available.

**Conclusion:** PANOPLY shows promise as a means to prioritize drugs based on clinical and multi-omics data for an individual cancer patient. Additional studies are needed to confirm this approach.

## INTRODUCTION

There has been substantial progress in the fight against cancer; however, cancer remains the second leading cause of death in the United States^1^. A major focus of cancer research has been the identification of oncogenic drivers and the development of drugs that selectively target those driver events. This approach has led to the development of agents that have been shown to successfully target the driver mutational events such as: trastuzumab, which targets *HER2+* breast cancer ^2,3^, imatinib, which inhibits the *BCR-ABLl* tyrosine kinase produced by the Philadelphia translocation in chronic myelogenous leukemia ^4^, vemurafenib for the treatment of *BRAF* V600E mutant malignant melanoma ^5^, agents targeting *EGFR* mutations in non-small cell lung carcinoma and ^6^, and crizotinib for non-small cell lung cancer with *ALK* rearrangements ^7^.

The mapping of the human genome has opened the door to the exploration of the tumor and environmental features to uncover the drivers of cancer and its resistance to treatment. A number of commercial gene sequencing platforms (e.g., Foundation One, Ambry Genetics) have been developed to identify tumor mutations used in clinical decision making. Most of these platforms are focused on detecting a limited number of gene abnormalities in specific genes and do not include comprehensive “multi-omics” data analysis. For many tumor types, this leads to the inability to link mutational drivers with druggable targets. A pressing need exists for better approaches to identify right drugs for an individual patient utilizing multi-omics data. While most of the studies utilize single omics data type to predict drug response, there are algorithms that use two or more genomic features to predict drug response in cancer cell lines ^8–10^and in The Cancer Genome Atlas (TCGA) subsets^11^. There are databases such as MD Anderson’s Personalized Cancer Therapy ^12^, Vanderbilt’s My Cancer Genome ^13^, the Broad Institute’s TARGET ^14^, TCGA ^15^, and the Catalogue of Somatic Mutations in Cancer (COSMIC) ^16^ containing information on the frequency of alterations in thousands of patients with cancer. Programs such as DriverNet ^17^, IntOGen ^18^, analyze a single type of omics data, such as somatic mutations, to identify potential driver genes. Other programs, such as XSeq^19^, OncoRep^20^, OncoIMPACT^21^, and iCAGES ^22^ integrate on somatic mutations and/or copy number alterations (CNAs), and/or gene expression. Although integrating these data types represents substantial progress toward the full molecular characterization needed for precision cancer care, no comprehensive method for integrating clinical and multi-omics data has yet been developed and validated for selecting the most compatible agents for a given patient’s -omics profile. Thus, we built a workflow called PANOPLY (**P**recision c**a**ncer ge**no**mic re**p**ort: single samp**l**e inventor**y**) that identifies molecular alterations that are unique to a cancer patient compared to matched-controls with similar disease characteristics who had a favorable clinical course and then performs a comprehensive, integrated multi-omics analysis to identify druggable genomic events for an individual’s tumor. The results are summarized in a report which the patient’s medical oncology team can use the data to choose the particular agent to be administered.

In brief, PANOPLY uses machine learning and knowledge-driven network analysis to analyze patient-specific alterations (CNA, germline and somatic alterations from DNA, and RNA gene expression and expressed mutations) driving oncogenesis and prioritizes drugs which target the networks and pathways associated with these cancer-driving alterations. We describe the workflow and provide examples using both institutional and publicly-available datasets where PANOPLY was used to identify drugs for individual patients and subgroups of patients whose disease is resistant to standard chemotherapy. We confirmed PANOPLY predictions in a patient with chemo-resistant triple-negative breast cancer (TNBC) using patient-derived xenografts (PDXs) by testing the top drugs for that patient.

## MATERIALS AND METHODS

### Data Sources

Datasets used in the manuscript are described in Data Sources section of the Supplementary Methods.

### PANOPLY workflow

The PANOPLY workflow is available for download as an R package from http://kalarikrlab.org/Software/Panoply.html and GitHub. A high-level overview of the workflow is shown in Figure 1A. Steps 1–4 are provided in the Supplementary Methods.

**Figure 1A).**
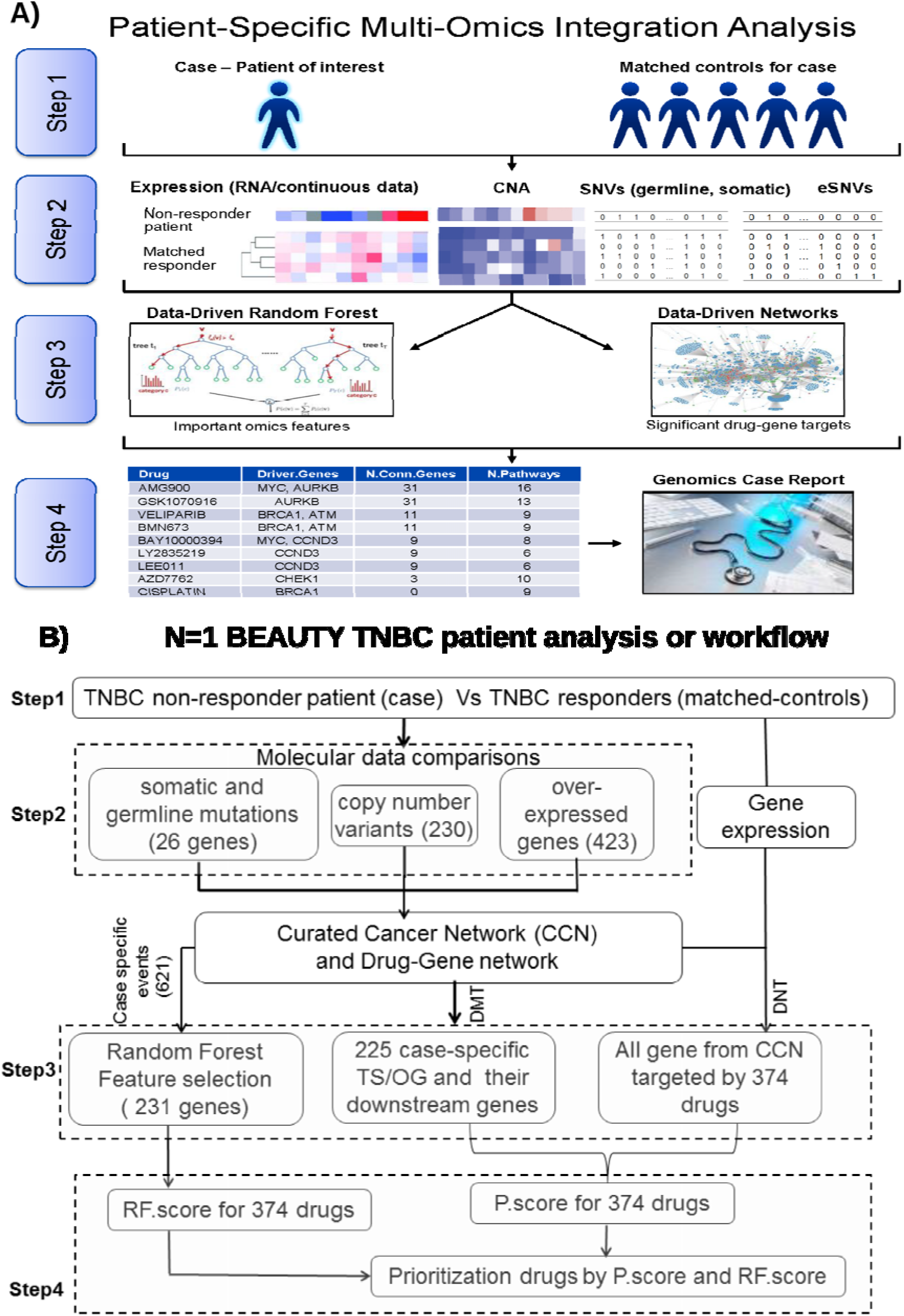
A high-level overview of PANOPLY. Step 1: A patient of interest is compared with matched-controls. Step 2: For each subject, gene expression data, copy number alteration (CNA), single nucleotide variant (SNV), and expressed single nucleotide variant (eSNV) data will be provided to identify case-specific driver alterations. Step 3: Multi-omics data will be provided to the random forest and network analysis methods to identify the top prioritized drugs to target genes that are driving oncogenesis in the patient. Step 4: A genomics case report listing the drugs to prioritize, based on their ability to target driver mutations and their dysregulated gene networks, is generated for researchers and clinicians. B) **An in-depth description of steps using a case example**. Application of PANOPLY to BEAUTY triple-negative breast cancer (TNBC) patient. Step 1 is a comparison of TNBC standard chemotherapy non-responder patient to a set a matched controls who have TNBC disease and have responded to standard chemotherapy. Step 2 is where all the molecular data comparisons are performed to identify case-specific events compared to matched-controls. In Step 3, random forest analysis, drug network test (DNT) and drug meta test (DMT) are conducted using the features obtained from Step 2. In Step 4, P.score (combination of network tests) and RF.score (random forest score) for anti-cancer drugs are calculated and prioritized based on high P.score and low RF.score.

### False Positive Rate (FPR) and True Positive Rate (TPR) Simulations

We evaluated FPR under three scenarios of varying levels of correlation between the drug-gene networks and the gene-gene networks: Sc1) all gene-gene and drug-gene networks; Sc2) reduced gene-gene networks, complete drug-gene networks; and Sc3) reduced gene-gene and reduced drug-gene networks. We describe the justification for these scenarios in the Supplementary Methods, where the binary clustering of the gene-gene and drug-gene networks are illustrated in Figures S2 and S3, respectively.

We evaluated the TPR of the Panoply workflow using just the simulated multivariate normal (MVN) data, by spiking in increased amounts of expression for a subset of genes in the Olaparib gene-drug network. Using the same framework as the FPR simulations, we changed the mean expression level for a subset of genes for the cases, but not the matched-controls (Supplementary Methods).

### Confirmation of PANOPLY’s Drug Predictions With PDXs

To validate PANOPLY’s drug predictions, we tested PDX models ^23^ obtained from BEAUTY ^24^ study (BC_051_1_1) for a TNBC patient whose tumor did not respond to neoadjuvant paclitaxel and anthracycline/cyclophosphamide treatment (Supplementary Methods).

## RESULTS

### Statistical Performance of PANOPLY Using Simulated Data

*FPR:* In Supplementary Figure S4 we observed the FPR for DNT, DMT, and P.score are controlled near the nominal α = 0.05 and 0.01 levels under a typical analysis with PANOPLY. We evaluated three scenarios (Sc1-Sc3) with varying correlation of the gene-gene and drug-gene networks and with varying matched-control set sizes M=2, 4, 8, and 16. Table 1 shows the results for M=8 for DNT and DMT for all scenarios with the two datasets. The FPRs are slightly higher in the TCGA breast cancer normal-adjacent samples than the simulated MVN set. The DNT error rates are adequate for all three simulations scenarios Sc1-Sc3, but perhaps too conservative in Sc2 and Sc3. The DMT was observed to be higher than the nominal levels in all scenarios Sc1-Sc3 in both datasets but is much closer to the nominal level when the correlation cancer gene networks are reduced in both Sc2 and Sc3. In summary, the suggested setting for running the workflow is scenario Sc2, where only the patient-specific events are considered for testing all 374 drugs.

**Table 1.** FPR and TPR rates for DNT and DMT test p-values at α = 0.05 using simulated multivariate normal (MVN) distribution (N=500 sets) and TCGA normal-adjacent data (N=94 sets), for matched control size M=8, and scenarios Sc1-Sc3. For TPR, we chose three subsets of genes that are targeted differently by the Olaparib drug-gene network: Set-A) the complete drug-gene network for Olaparib; Set-B) a small network of BRCA2-specific genes, and Set-C) a mixture of genes that are sub-networks of the ATM/ATR networks, which are larger than the BRCA2 network. Gene sets A, B, and C simulated with mean expression for the case as 2 and 3 gene-specific standard deviations (sd) higher than the controls.

| Scenario |  |  | Test | False Positive Rate |  | True Positive Rate |  |  |  |  |  |
| --- | --- | --- | --- | --- | --- | --- | --- | --- | --- | --- | --- |
|  | Drivers | Drugs |  | Null |  | Set A |  | Set B |  | Set C |  |
|  |  |  |  | MVN | TCGA | MVN (sd 2) | MVN (sd 3) | MVN (sd 2) | MVN (sd 3) | MVN (sd 2) | MVN (sd 3) |
| Sc1: All Driver Genes and Drugs | 429 | 374 | DNT | 0.029 | 0.04 | 0.582 | 0.926 | 0.030 | 0.062 | 0.108 | 0.294 |
|  |  |  | DMT | 0.231 | 0.208 | 0.522 | 0.588 | 0.496 | 0.542 | 0.498 | 0.544 |
| Sc2: Patient Observed Driver Genes, all Drugs | 120 | 374 | DNT | 0.031 | 0.038 | 0.612 | 0.942 | 0.038 | 0.082 | 0.132 | 0.286 |
|  |  |  | DMT | 0.065 | 0.104 | 0.390 | 0.410 | 0.320 | 0.326 | 0.528 | 0.590 |
| Sc3: Reduced Correlation Drivers and Drugs | 175 | 117 | DNT | 0.008 | 0.01 |  |  |  |  |  |  |
|  |  |  | DMT | 0.08 | 0.088 |  |  |  |  |  |  |

*TPR:* We quantify TPR as the proportion of simulated datasets for which the DNT and DMT p-values for Olaparib are significant in two ways: the p-value is less than α = 0.05, and the p-value is one of the ten lowest of all drugs. As shown in Table 1 for M=8 and α = 0.05, the TPR for DNT and DMT are comparable with Gene Set-A while DNT TPR dramatically drops off with more realistic network-specific gene-gene networks that are activated (Gene Sets B and C), as expected. Complete results for DNT, DMT, and P.score are discussed in the Supplementary Material. Supplementary Figures S5 and S6 show the TPR is robust to the size of the reference sample population (M=2, 4, 8, and 16), the top-10 metric verifies the α = 0.05 TPR rates, and that P.score is useful in ranking drugs that perform well across DNT or DMT. Furthermore, Supplementary Figure S7 shows the TPR gains in DMT and P.score for incremental increases in overexpression of genes targeted by a drug; we also observed higher TPR values in the model when DNA aberration events are spiked in addition to RNA expression (Supplementary Results).

### N=1 Case Study

We applied PANOPLY workflow to prioritize drugs for a BEAUTY patient (case BC_051_1_1) who did not respond to neoadjuvant chemotherapy for whom a set of matched controls (n=9) were found among the BEAUTY TNBC patients who had a pathologic complete response (pCR) to neoadjuvant chemotherapy. The PANOPLY report for that patient is available in the Supplementary (file-Supplementary_PANOPLY_BEAUTY_Patient_Report.pdf). Below we have (1) the experimental validation of a PANOPLY-prioritized drug using PDX models and (2) comparison of PANOPLY analysis for a BC_051_1 patient with other methods.

1. *Experiment validation:* Somatic, germline mutations, CNA, gene expression, and expressed mutation data for the case was compared and contrasted with her matched controls using PANOPLY workflow. (Resulting tables for case: BC_051_1_1 are discussed in detail in the Supplementary Results). A high-level step by step analysis of PANOPLY algorithm for this patient is shown in Figure 1B. The PANOPLY results indicated that olaparib was the most promising drug for this patient (Table 2). Figure 2A shows histologic images of the case’s tumor and a corresponding PDX, both from the pre-NAC and post-NAC time points. Pre- and post-NAC patient tumor, and its corresponding PDX had similar morphologic features and a triple negative staining pattern (Figure 2A). For both the pre-NAC and post-NAC PDXs, tumor volume at day 12 was significantly lower in the olaparib group than in the vehicle group (Wilcoxon rank sum test *P*=0.04 and *p* <0.001 respectively; Figure 2B). The PDX results show promise that this approach may be successful in identifying an effective therapy for patients.
2. *Comparison of PANOPLY with existing methods:* Recent bioinformatics software, such as iCages and oncorep, attempt to incorporate a tabulation of anti-cancer drug options targeting observed driver genes. These softwares were developed independently and with their own design assumptions and intent. Table 3 presents a summary of these two software implementations, in comparison to Panoply. We were able to generate a similar iCages report with default settings for case BC_051_1_1, by providing required somatic mutation (VCF format) and the copy number alteration data (BED format). We were not able to configure the current architecture of the Amazon web services, required for the omics_pipe workflow (which precedes the oncorep analysis module).

**Figure 2.**
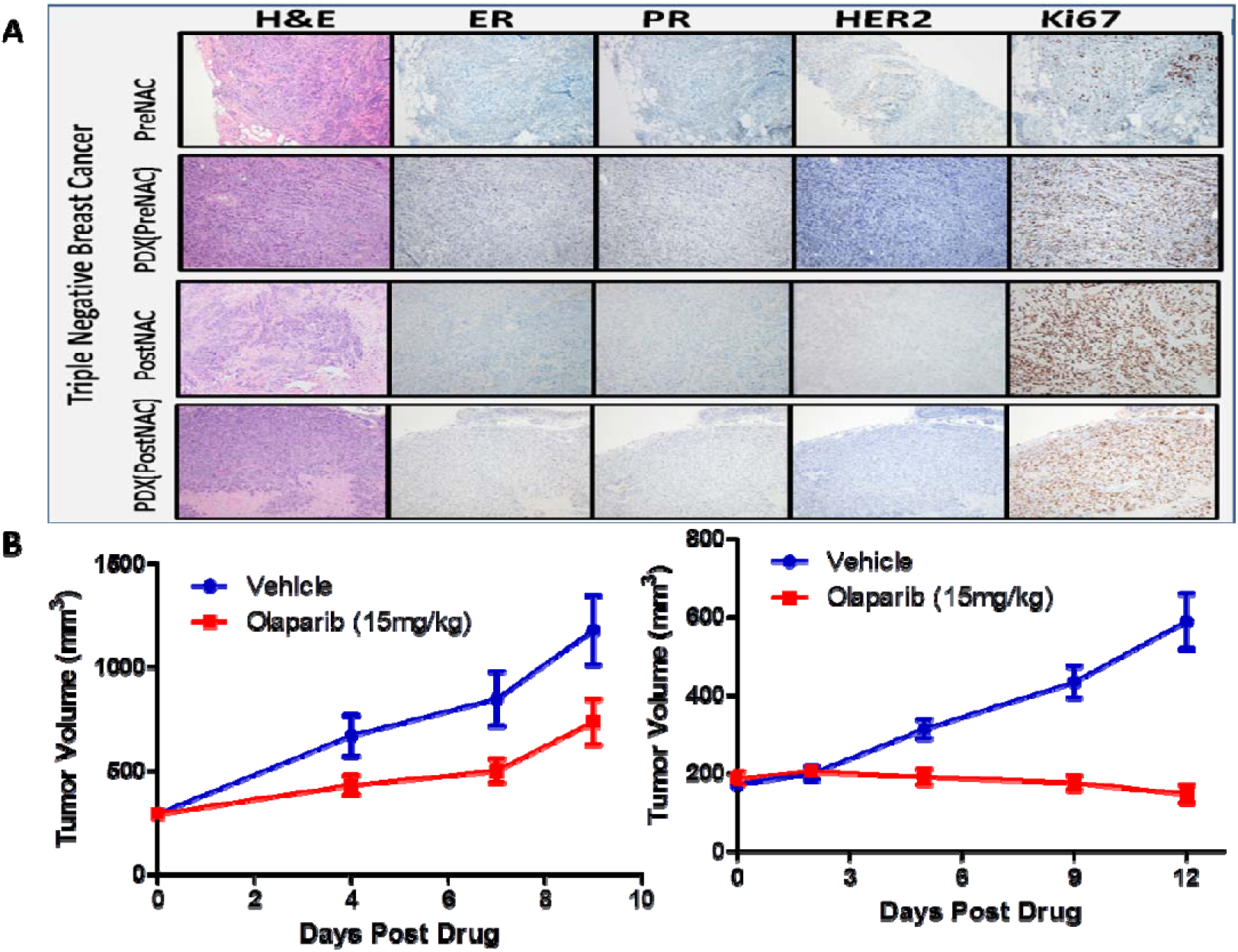
Patient-derived xenografts (PDXs) confirm PANOPLY’s prediction that olaparib is an effective treatment for a patient with chemoresistant TNBC (BC_051_1_1). A) The top panel shows histological stains of the patient’s tumor and PDXs (pre or post-treatment). B) Cytotoxicity data shows the PDXs response to the top predicted drug olaparib compared to the no treatment (the left plot shows the olaparib drug response data from pre-treatment mice, whereas the data from the right shows the data from post-treatment PDX models. Both the datasets were generated using the Vehicle as controls).

**Table 2:** Top drugs recommended by PANOPLY for BEAUTY patient (BC_051_1_1) with chemoresistant triple-negative breast cancer and PDX model. PANOPLY identified pathways associated with defective DNA repair for BC_051_1_1 patient. However, the top four drugs identified and prioritized by PANPOLY were all PARP inhibitors and included olaparib, talazoparib, rucaparib, and veliparib. While any one of these PARP inhibitors could have been chosen for in vivo studies, we focused on olaparib, given its high ranking and, given that it was the only PARP FDA approved at the time. Of note, there were other drugs prioritized lower but still known to demonstrate antitumor activity in the presence of DNA repair deficiency, including carboplatin and CHEK inhibitors.

| **Drug** | **Case Driver Genes<sup>a</sup>** | **Number of Pathways<sup>b</sup>** | **P.score** | **RF.score** |
| --- | --- | --- | --- | --- |
| OLAPARIB | ATR,ATM,BRCA2,BRCA1 | 43 | 3.009632 | 0.003743 |
| BMN673 | ATR,ATM,BRCA2,BRCA1 | 32 | 2.70456 | 0.004679 |
| RUCAPARIB | ATR,ATM,BRCA2,BRCA1 | 32 | 2.7046 | 0.0047 |
| VELIPARIB | ATR,ATM,BRCA2,BRCA1 | 32 | 2.7046 | 0.0047 |
| AMG900 | AURKB,AURKA | 8 | 2.610079 | 0.001857 |
| AZD7762 | CHEK1,CHEK2 | 13 | 2.566379 | 0.001786 |

<sup>a</sup> Genes on PANOPLY's cancer gene list that are targeted by FDA-approved drugs
<sup>b</sup> Number of pathways that may be affected, based on drug-gene target interactions Clinical Characteristics of this cancer patient can be found in the Table 1 of Supplementary\_PANOPLY\_BEAUTY\_Patient\_Report.pdf

**Table 3:**
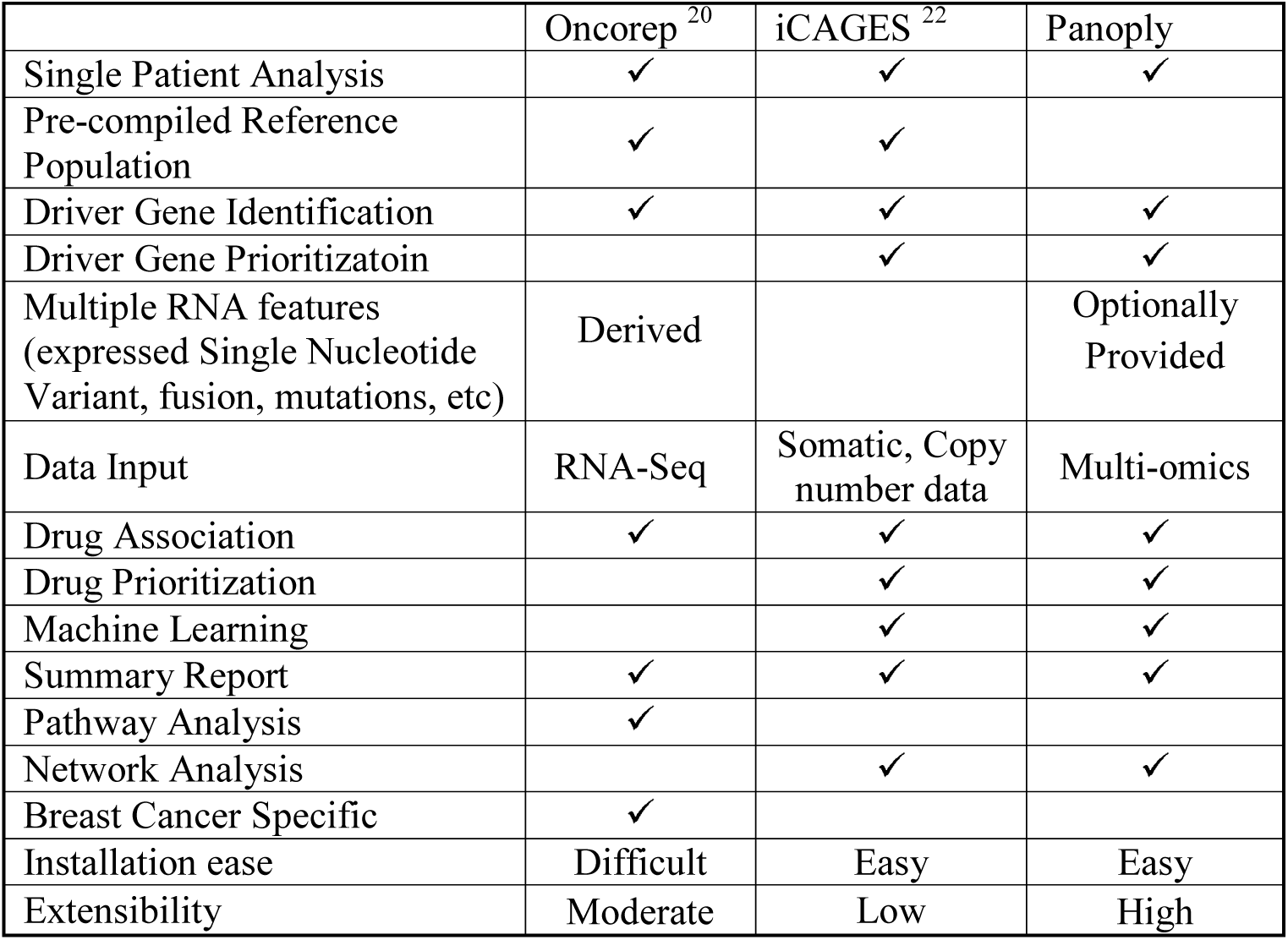
Comparison of oncorep ^20^, iCAGES ^22^ and Panoply methods

### Extensibility of the Workflow to Cohort Studies and Public Domain Datasets

*PANOPLY Drug Predictions for Patients with Chemo-resistant, TNBC cohort:* Panoply provides a prioritized drug list for each patient in the cohort. This list corresponds to a unique set of gene targets for each patient, which can be compared and contrasted with similar chemo-resistant patients using existing clustering methods. The genomic characteristics of these clusters can be reverse engineered to find qualifying genomic events which would qualify future patients for drug ‘bucket’ trials. An illustration of this application of PANAPOLY is provided using the 17 BEAUTY patients with chemoresistant TNBC. NMF clustering ^25^ was performed with the drug priority scores of these 17 cases. Based on the cophenetic and average silhouette scores, two clusters were selected to be optimal. The percentile ranking of top 10% (35/344) drugs was aggregated per sample cluster using the median score and presented as a heatmap (Figure 3A). The target genes of the drug clusters were collated, and a word cloud was generated with the targets (Figure 3B). As shown in Figure 3A, the cluster 1 consists of nine samples; the patients in that group primarily consist of kinase-inhibitors as their top prioritized drugs (drugs=16) and the drugs in that cluster can predominantly target the PIK3CA-mTOR-AKT signaling pathways. The other cluster from Figure 3A consists of a set of prioritized drugs (drugs=19) for eight patients, as shown in Figure 3B these drugs can primarily target genes associated with cell cycle control, specifically targeting the histone deacetylases *(HDAC1)* and the Aurora kinases A and B.

**Figure 3.**
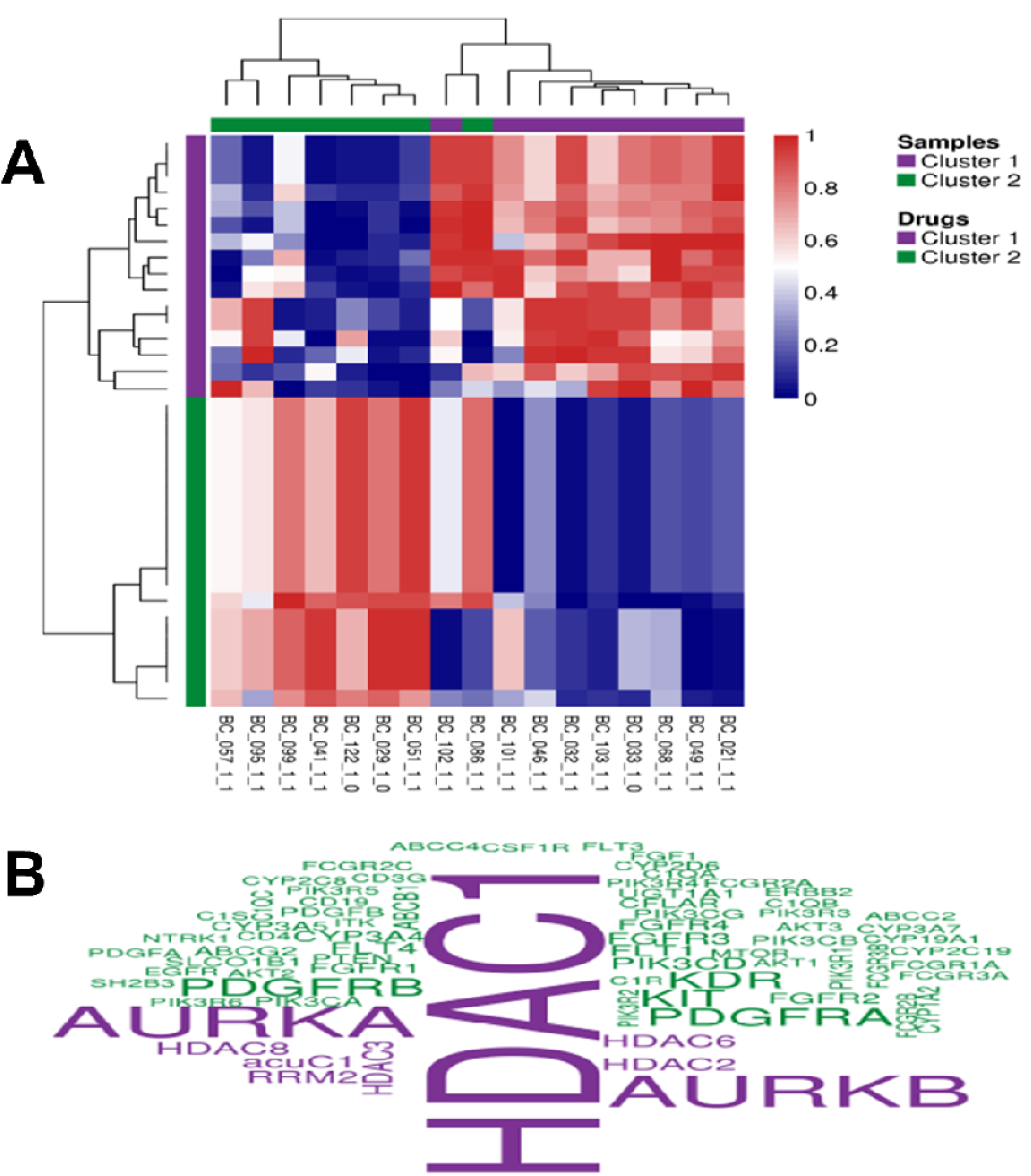
Clustering and word cloud plots of 17 TNBC chemotherapy-resistant patients. A) Two-way hierarchical clustering of the top 10% of the drugs predicted by PANOPLY for 17 basal TNBC patients. The heatmap shows that there are two sample and drug clusters are implicated in the NMF clustering analysis. B) Word cloud of the target genes from the two drug clusters predicted by the NMF analysis.

*PANOPLY drug predictions for public dataset such as TCGA with molecular and clinical data:* In here, we present the capability of PANOPLY workflow to be executed with publically available datasets. An example COAD patient’s (TCGA-AA-3488) report is discussed briefly below and can be obtained from Supplementary_PANOPLY_TCGA_COAD_Patient_Report.pdf. The case displayed 645 driver genes: 226 CNAs not present in the case; in addition, 419 genes were over-expressed in the case tumor sample relative to the matched-control samples. Of the 226 CNAs, 113 were amplifications, and 113 were deletions. Of those BRAF, CDKN2A, CDKN2B, FGF10, IL7R, INHBA, JAK2, KEL, MAFA, NTRK1, NTRK2, PIK3CA, PRSS1, RBL1, SKIL, SMO, SOX2 and SPTA1 cancer-related genes were both amplified and overexpressed. Of the case’s 645 events, 53 genes are differentially expressed between the case and matched-controls and can be targeted by antineoplastic drugs. Based on the network and random forest analysis of driver genes and gene expression data, PANOPLY ranked the following drugs lestaurtinib (JAK2, NTRK1, NTRK2), LY2784544 (JAK2), GDC-0032(PIK3CA), NVP-BGT226 (MTOR, PIK3CA), regorafenib (MAPK11, RAF1, BRAF, KRAS, KIT, FGFR1/2,PDGFRA/B, ARAF, KDR, EPHA2, ABL1, NTRK1, CYP3A4, CYP2C8/9, CYP2B6, CYP2C19,ABCB1, ABCG2, UGT1A1/9,FLT1/4, RET, TEK and DDR2) and ARQ736 (BRAF) as significant for patient TCGA-AA-3488 with significant P.score and RF.score (Table **S1**).

Similarly, we have also applied PANOPLY to TCGA breast cancer data (BRCA), and the exemplary report was prepared for a TCGA tumor (TCGA-AR-A1AR) sample and is presented in the Supplementary.

## DISCUSSION

In creating PANOPLY, our goal was to develop a flexible workflow capable of analyzing multiple forms of -omics and clinical data to identify driver genes, their effects on gene networks and the drugs capable of targeting altered gene networks in cancer patients. Using gene expression data, CNA, and DNA variants from publicly available and in-house datasets, we demonstrate that PANOPLY holds promise in identifying agents capable of targeting driver gene-induced changes, both for individual patients and for subgroups of patients that share a cancer subtype and response pattern. This is evident in the example where PANOPLY’s prioritization of olaparib as a promising treatment for a patient with chemoresistant TNBC and that agent was found to reduce tumor size when applied to that patient’s xenografts. When the same patient’s data was run through iCages, the drug with the highest iCageDrugScore (0.52) was Doxorubicin, the drug administered as part of the patient’s NAC regimen that failed produce a pCR. While Olaparib scored (1.3 × 10^−4^) much lower among potential agents. Doxorubicin intercalates DNA and thereby indirectly targets TP53 (TOP2A), which is observed in a substantial proportion of cancer patients. Co-considering associated genes involved with DNA damage repair and/or active cellular uptake would presumably provide a more accurate prediction of drug efficacy, as implemented by Panoply.

The PANOPLY workflow currently analyzes the patient's molecular and clinical data together along with the knowledge databases such as Reactome, DGI-db, and others for drug-gene network analysis. This represents a substantial advancement relative to existing programs, such as XSeq ^19^, OncoRep ^20^, and OncoIMPACT ^21^, which integrate only molecular data such as somatic mutations or CNAs, and gene expression. Currently, medical oncologists have access to genomics reports that were generated using a limited target panel for decision making. Working closely with clinicians, basic scientists, and pharmacologists, we have developed PANOPLY to integrate molecular, clinical, and drug data to prioritize targets and facilitate individualized treatment for the patients. Using clinical and molecular profiles of the patient’s disease, PANOPLY provides a personalized list of prioritized drugs along with links to literature concerning drug efficacy. The oncologist will still have to go through the list and refine drug selection based on the inherent clinical knowledge, ancillary clinical trials, and insurance coverage availability.

Obtaining the whole genome and transcriptome data is challenging for implementation of PANOPLY in a routine setting. PANOPLY’s framework can be customized to analyze focused gene panels with expression and mutation data by modifying the gene adjacency matrix (Reactome.adj module) as described in the user manual of the PANOPLY package. However, we do not recommend such practice since it is going to be a compromise of our algorithm and its capabilities. There are several studies ^24,26–32^ that have both molecular and clinical data publically available; however, most of these analyses use a cohort-based comparison that may not be specific to an individual. As individual interventions become a reality on a routine basis in the precision medicine era, PANOPLY’s framework to harness the current knowledge in genomics and guide patient treatment based on clinical and multi-omics data becomes more valuable.

Presently, generating multi-omics data for a patient is still cost-prohibiting but as the cost of clinical sequencing continues to decline ^33–36^ it may become less of a concern and more feasible in the near future. Obtaining clinical data and designing matched-control set is a laborious task. However, extensive application of the electronic medical records in clinics and the recent development of functions like “optmatch” in R would provide ways for the availability of optimally matched cohorts. For now, if no matched-controls are available for a study, we recommend using TCGA multi-omics and clinical data to design matched-control set. Although the performance might be compromised due to the batch effects between data sets, appropriate adjustments of data should minimize the effect. As an option, we provide a module in PANOPLY to perform adjustment to counter for the batch effect. Another limitation of PANOPLY is that the method cannot delineate the clinical effectiveness of a similar class of drugs. We plan to accomplish this in future by bringing in drug knowledge such as chemical structure, molecular size, and drug dosage. Our method is dependent on drug-gene target annotations that are heavily biased by product literature and databases. Moreover, clinical translation of PANOPLY results is constrained by the cost-effectiveness in developing PDXs for drug validation.

In summary, PANOPLY is a flexible framework that can integrate many other types of -omics data, including protein expression, methylation expression, structural variants, circular RNAs, long non-coding RNAs, and fusion data with modifications to its code. PANOPLY’s framework can also be extended to the metastatic setting; additional clinical data such as prior exposure to drugs, features of primary and recurrence disease would be required. Additional genomics data sets are needed to modify the existing approach for metastatic tumors. As more is learned about the molecular underpinning of cancer using various resources, we plan to expand our knowledge base to improve PANOPLY’s predictions using PDX and cell-lines. We have validated PANOPLY’s predictions in a single patient at present using PDXs, in future, we plan to validate PANOPLY-predicted drugs in PDXs derived from additional patients. Like several other groups and we have shown, PDX models faithfully represent tumor biology^23,37^, so these results should provide insight into PANOPLY’s reliability. In conclusion, our results indicate that combining multiple sources of -omics and clinical data to predict promising agents for a patient or groups of patients with cancer is feasible. With further validations, PANOPLY can be a tool to help clinicians in their decision-making process.

## ACKNOWLEDGEMENTS

KRK, JPS, KJT, XT, EEC, JY, PTV, JNI, RMW, LW, JCB, MPG, and VJS are funded in part by the Mayo Clinic Center for Individualized Medicine (CIM). KRK, RMW, LW, MPG, JNI, and VJS are funded in part by the Mayo Clinic Breast Specialized Program of Research Excellence (SPORE) (P50CA116201). KRK, RMV, LW, MPG and JCB are funded in part by the U54 GM114838. MPG and VJS are funded in part by the Mayo Comprehensive Cancer Center Grant (P30CA 15083-43). KRK, JS, KJT, XT, EEC, and PTV are funded by the Division of Biostatistics and Informatics at the Mayo Clinic.

## REFERENCES

1. CDC: Leading causes of death, 2015

2. Perez EA, Reinholz MM, Hillman DW, et al: HER2 and chromosome 17 effect on patient outcome in the N9831 adjuvant trastuzumab trial. Journal of clinical oncology: official journal of the American Society of Clinical Oncology 28:4307–15, 2010

3. Perez EA, Dueck AC, McCullough AE, et al: Predictability of adjuvant trastuzumab benefit in N9831 patients using the ASCO/CAP HER2-positivity criteria. Journal of the National Cancer Institute 104:159–62, 2012

4. Gambacorti-Passerini C, Antolini L, Mahon F, et al: Multicenter independent assessment of outcomes in chronic myeloid leukemia patients treated with imatinib. J Natl Cancer Inst 103:553–61, 2011

5. Chapman P, Hauschild A, Robert C, et al: Improved survival with vemurafenib in melanoma with BRAF V600E mutation. N Engl J Med 364:2507–16, 2011

6. Mok T, Wu Y, Thongprasert S, et al: Gefitinib or caboplatin-paclitaxel in pulmonary adenocarcinoma N Engl J Med 361:947–57, 2009

7. Kwak E, Bang Y, Camidge D, et al: Anaplastic lymphoma kinase inhibition in non-small-cell lung cancer. N Engl J Med 363:1693–703, 2010

8. Dong Z, Zhang N, Li C, et al: Anticancer drug sensitivity prediction in cell lines from baseline gene expression through recursive feature selection. BMC cancer 15:489, 2015

9. Zhang N, Wang H, Fang Y, et al: Predicting Anticancer Drug Responses Using a Dual-Layer Integrated Cell Line-Drug Network Model. PLoS computational biology 11:e1004498, 2015

10. Qin Y, Conley AP, Grimm EA, et al: A tool for discovering drug sensitivity and gene expression associations in cancer cells. PloS one 12:e0176763, 2017

11. Geeleher P, Cox NJ, Huang RS: Cancer biomarker discovery is improved by accounting for variability in general levels of drug sensitivity in pre-clinical models. Genome biology 17:190, 2016

12. Meric-Bernstam F, Johnson A, Holla V, et al: A decision support framework for genomically informed investigational cancer therapy. Journal of the National Cancer Institute 107, 2015

13. Yeh P, Chen H, Andrews J, et al: DNA-Mutation Inventory to Refine and Enhance Cancer Treatment (DIRECT): a catalog of clinically relevant cancer mutations to enable genome-directed anticancer therapy. Clinical cancer research: an official journal of the American Association for Cancer Research 19:1894–901, 2013

14. Van Allen EM, Wagle N, Stojanov P, et al: Whole-exome sequencing and clinical interpretation of formalin-fixed, paraffin-embedded tumor samples to guide precision cancer medicine. Nature medicine 20:682–8, 2014

15. Cerami E, Gao J, Dogrusoz U, et al: The cBio cancer genomics portal: an open platform for exploring multidimensional cancer genomics data. Cancer discovery 2:401–4, 2012

16. Forbes SA, Beare D, Gunasekaran P, et al: COSMIC: exploring the world’s knowledge of somatic mutations in human cancer. Nucleic acids research 43:D805–11, 2015

17. Bashashati A, Haffari G, Ding J, et al: DriverNet: uncovering the impact of somatic driver mutations on transcriptional networks in cancer. Genome biology 13:R124, 2012

18. Gundem G, Perez-Llamas C, Jene-Sanz A, et al: IntOGen: integration and data mining of multidimensional oncogenomic data. Nature methods 7:92–3, 2010

19. Ding J, McConechy MK, Horlings HM, et al: Systematic analysis of somatic mutations impacting gene expression in 12 tumour types. Nature communications 6:8554, 2015

20. Meissner T, Fisch KM, Gioia L, et al: OncoRep: an n-of-1 reporting tool to support genome-guided treatment for breast cancer patients using RNA-sequencing. BMC medical genomics 8:24, 2015

21. Bertrand D, Chng KR, Sherbaf FG, et al: Patient-specific driver gene prediction and risk assessment through integrated network analysis of cancer omics profiles. Nucleic acids research 43:e44, 2015

22. Dong C, Guo Y, Yang H, et al: iCAGES: integrated CAncer GEnome Score for comprehensively prioritizing driver genes in personal cancer genomes. Genome medicine 8:135, 2016

23. Yu J, Qin B, Moyer AM, et al: Establishing and characterizing patient-derived xenografts using pre-chemotherapy percutaneous biopsy and post-chemotherapy surgical samples from a prospective neoadjuvant breast cancer study. Breast cancer research: BCR 19:130, 2017

24. Goetz MP, Kalari KR, Suman VJ, et al: Tumor Sequencing and Patient-Derived Xenografts in the Neoadjuvant Treatment of Breast Cancer. Journal of the National Cancer Institute 109, 2017

25. Mirzal A: Nonparametric Tikhonov Regularized NMF and Its Application in Cancer Clustering. IEEE/ACM transactions on computational biology and bioinformatics 11:1208–17, 2014

26. Kong J, Cooper LA, Wang F, et al: Integrative, multimodal analysis of glioblastoma using TCGA molecular data, pathology images, and clinical outcomes. IEEE transactions on bio-medical engineering 58:3469–74, 2011

27. Brodie SA, Li G, Brandes JC: Molecular characteristics of non-small cell lung cancer with reduced CHFR expression in The Cancer Genome Atlas (TCGA) project. Respiratory medicine 109:131–6, 2015

28. Zhang Q, Burdette JE, Wang JP: Integrative network analysis of TCGA data for ovarian cancer. Bmc Systems Biology 8:1338, 2014

29. Neapolitan R, Horvath CM, Jiang X: Pan-cancer analysis of TCGA data reveals notable signaling pathways. BMC cancer 15:516, 2015

30. Huang Z, Duan H, Li H: Identification of Gene Expression Pattern Related to Breast Cancer Survival Using Integrated TCGA Datasets and Genomic Tools. BioMed research international 2015:878546, 2015

31. Sun H, Yan L, Tu R, et al: Expression Profiles of Endometrial Carcinoma by Integrative Analysis of TCGA Data. Gynecologic and obstetric investigation 82:30–38, 2017

32. Spainhour JC, Qiu P: Identification of gene-drug interactions that impact patient survival in TCGA. BMC bioinformatics 17:409, 2016

33. Mardis ER: Anticipating the 1,000 dollar genome. Genome biology 7:112, 2006

34. Singh S: The hundred-dollar genome: a health care cart before the genomic horse. CMAJ: Canadian Medical Association journal = journal de l’Association medicale canadienne 190:E514, 2018

35. Morrison C, Trump D, Nowak JA: How Will the “$1,000 Dollar Genome“ Meet Reality (and Centers for Medicare & Medicaid Services)? Archives of pathology & laboratory medicine 139:581–2, 2015

36. Selwood DL: Beyond the hundred dollar genome-drug discovery futures. Chemical biology & drug design 81:1–4, 2013

37. Hou J, Wang L: FKBP5 as a selection biomarker for gemcitabine and Akt inhibitors in treatment of pancreatic cancer. PloS one 7:e36252, 2012

